# Explaining the rapid evolution of mammalian meiotic recombination proteins

**DOI:** 10.64898/2026.06.01.729254

**Authors:** Julien Joseph, Paola Montoya, Frédéric Baudat, Aurora Ruiz-Herrera, Julien Clavel

## Abstract

Meiotic recombination — the exchange of genetic material between parental chromosomes during gamete production — is critical for fertility, genome stability, and evolutionary adaptation. In eukaryotes, meiotic recombination is carried out by a deeply conserved molecular machinery. Despite this conservation, the sequences of proteins involved in meiotic recombination evolve at a remarkably high rate in mammalian species. Several biological processes have been proposed to explain this rapid evolution, but none have been quantitatively tested. In this study, we analyzed the variation in evolutionary rates of 100 recombination proteins across approximately 400 placental mammals. Our results show that this rapid evolution is primarily driven by lower levels of purifying selection compared to the rest of the proteome. We show that, although selective pressures exerted on recombination proteins are generally shared across mammals, a few recombination proteins exhibit strong variation in selective pressures. Contrary to previous hypotheses, we demonstrate that chromosome number and genome-wide recombination rates do not account for much of this variation. Instead, these variations are primarily associated with chromosome pairing and synapsis proteins, which tend to experience increased selective pressures throughout mammalian evolution, especially in large and long-lived species. This pattern probably reflects more intense selection for stability of chromosome pairing proteins in the oocytes of long-lived species, in which chromosomes sometimes need to stay paired for decades.

## Introduction

During meiosis — the production of sex cells — parental chromosomes exchange genetic material through homologous recombination. Meiotic recombination begins with the formation of hundreds of programmed double-stranded breaks (DSBs) throughout the genome. These breaks are then repaired using the homologous chromosome as a template. The repair process involves either an exchange of a chromosome arm, referred to as a crossover (CO), or a transfer of genetic material from the intact chromosome to the broken one, referred to as a non-crossover (NCO). This exchange of genetic information increases genetic shuffling, thereby enhancing genetic and functional diversity in the population (Hill and Robertson, 1966; Smith and Haigh, 1973; Felsenstein, 1974; Charlesworth et al., 1993). Moreover, in most eukaryotes, meiotic recombination is intrinsically linked to homologous chromosome pairing and synapsis and is essential to transmit chromosomes to the next generation during sexual reproduction (Zickler and Kleckner, 2023). However, meiotic recombination can be highly mutagenic (Magni and Von Borstel, 1962; Magni, 1963; Rattray et al., 2015; Pratto et al., 2014; Halldorsson et al., 2019; Arbel-Eden and Simchen, 2019; Hinch et al., 2023) and can lead to transmission distortion through biased gene conversion (Boulton et al., 1997; Duret and Galtier, 2009). Therefore, it is critical to understand how meiotic recombination is regulated in different species in order to understand their evolution. Interestingly, despite variability in both the meiotic recombination rates and the distribution of recombination events across the genome (Segura et al., 2013; Priore and Pigozzi, 2017; Stapley et al., 2017; Wang et al., 2019), major recombination steps are strikingly conserved across eukaryotes. (reviewed in Marín-Gual et al. (2022); Arter and Keeney (2023); Marin-Gual et al. (2025)). Moreover, proteins involved in different steps of meiotic recombination are often homologous in distantly related eukaryotes, often performing the same function (Arter and Keeney, 2023). However, these homologous proteins exhibit strong sequence variability, with proteins that are nearly unrecognizable between plants, animals, and fungi (Arter and Keeney, 2023). This variability is likely due to the fact that these proteins evolve quickly compared to the rest of the genome over short time scales, as observed in mammals (Dapper and Payseur, 2019; Arter et al., 2025), birds (Szasz-Green et al., 2025), maize (Sidhu et al., 2017) and *Drosophila* (Anderson et al., 2009; Brand et al., 2018; Zakerzade et al., 2025). The reasons for this rapid evolution are still not well understood, but could shed light on the biological processes influencing the evolution of meiosis itself. Several hypotheses have been proposed so far.

A non-adaptive hypothesis states that, because many recombination proteins are involved in a single process, those proteins are subject to lower selective pressures than the rest of the proteome (Dapper and Payseur, 2019). Therefore, mutations in these proteins are less often pleiotropic and have fewer deleterious effects. This is particularly true for recombination proteins that are only expressed during meiosis, but less so for proteins that are also involved in homologous repair of spontaneous double stranded breaks in somatic tissues. Under this hypothesis, the rapid evolution of recombination proteins is due to more relaxed purifying selection compared to the rest of the proteome. In contrast, most adaptive hypotheses involve recurrent selective pressures that change the recombination rate (Arter and Keeney, 2023). These pressures can be caused by environmental changes that favor modifiers alleles increasing the recombination rate because individuals carrying these alleles are more likely to generate combinations of loci that increase fitness in stressful conditions (Otto and Barton, 1997). Additionally, a recent study demonstrated that a modifier of the recombination landscape can be selected in one environment but not in others, potentially leading to recurrent selection on proteins that act on the recombination landscape (Parée et al., 2025). Chromosome number and structure also impose constraints on the evolution of recombination rates (Capilla et al., 2016). Since one crossing-over (CO) per chromosome is necessary for proper chromosome segregation (Kaback et al., 1992), fissions and fusions of chromosomes will result in changes to the total number of COs (Pardo-Manuel de Villena and Sapienza, 2001; Fernandes et al., 2018; Vara et al., 2021; Brazier and Glémin, 2022; Marín-García et al., 2024). Recent evidence from natural populations demonstrate that chromosomal fusions in mice lead to a redistribution of COs towards telomeric regions in metacentric chromosomes (Marín-García et al., 2024). This repatterning is associated with increased levels of CO interference, reduced population-level recombination estimates, and pronounced genomic divergence. Recurrent chromosome fission and fusions can therefore change the extent to which recombination is needed, which can increase or decrease selective pressures on meiotic recombination proteins. The high incidence of positive selection inferred in meiotic recombination proteins in mammals (Dapper and Payseur, 2019; Arter et al., 2025), birds (Szasz-Green et al., 2025), *Drosophila* (Anderson et al., 2009) and yeasts (Arter et al., 2025) provides support for these adaptive hypotheses. Furthermore, some of these proteins have been shown to be associated with variation in recombination rates and distribution within and between populations (reviewed in Johnston (2024) and Payseur (2025)).

While these observations are insightful, they do not constitute a proper test of the hypotheses presented above. In particular, they do not provide a quantitative estimate of the extent to which positive selection for changes in genome-wide recombination rates can account for the rapid evolution of these proteins. First positive selection is very often overestimated in the presence of residual alignment errors (Jordan and Goldman, 2012; Selberg et al., 2025) or biased gene conversion (Galtier and Duret, 2007; Ratnakumar et al., 2010), and underestimated in selectively constrained proteins (Rodrigue and Lartillot, 2017). Second, positive selection can occur in any gene because of processes unrelated to their functions (Hartl and Taubes, 1996; Jones et al., 2017; Latrille et al., 2024; Joseph, 2024; Riffis et al., 2025). Third, many recombination proteins are involved in other cellular processes outside of meiosis (e.g. DNA repair, expression regulation), and positive selection in these proteins may be totally unrelated to meiosis. Finally, changes in recombination proteins can reflect changes of the recombination process beyond recombination rate and distribution. Indeed, other properties of meiotic recombination, such as speed, timing, accuracy, and fidelity, vary between species (Adler, 1996; Soares et al., 2009; Murat et al., 2023) and can significantly impact fertility and fitness (Mihola et al., 2021).

In this study, we aim to answer two questions by taking placental mammals as a case study. First, what is the relative contribution of positive and relaxed purifying selection to the high evolutionary rate of recombination proteins? Second, which ecological, demographic or cellular processes have the greatest impact on the rate at which these proteins evolve?

To address these questions, we investigated the evolutionary rates of 100 recombination and 100 randomly selected proteins across amino-acids and across species in 413 placental mammals. First, we confirm that the incidence of positive selection is slightly higher in recombination proteins compared to randomly selected proteins. However, their evolutionary rate is largely driven by low selective pressures, more strikingly so for meiosis-specific proteins. Second, we find that selective pressures acting on meiotic recombination proteins are broadly conserved across mammals, although a subset of proteins exhibits notable heterogeneity. Interestingly, we demonstrate that neither genome-wide recombination rates nor chromosome number account for this variation. Instead, it is better explained by functional changes in the way homologous chromosomes are paired and synapsed in large and long-lived mammals.

## Results

### The respective contribution of low purifying selection and recurrent adaptation

First, we evaluated the extent to which positive selection explains the rapid evolution of recombination proteins. For this, we used a classical codon model (Muse and Gaut, 1994) to compute the non-synonymous over synonymous substitution rate (*dN/dS*) for each site of 100 proteins with experimental evidence for a role in meiotic recombination, and 100 control proteins randomly selected across the genome. For each site, if the *dN/dS* is higher than one, it means that non-synonymous mutations fix at a higher rate than synonymous mutations, expected to be neutral (Miyata and Yasunaga, 1980; Nei and Gojobori, 1986). This suggest that this site has been undergoing one or several episodes of positive selection. One caveat with this approach is that its power is drastically reduced in sites that are generally under purifying selection, which correspond to the majority of sites in protein-coding genes (Rodrigue and Lartillot, 2017; Latrille et al., 2023). Therefore, using this approach will tend to underestimate positive selection in the proteins that are more constrained (Rodrigue and Lartillot, 2017). This compromises our capacity to disentangle the contribution of the strength of purifying selection and the rate of positive selection to the *dN/dS* of a protein.

A simple alternative is to compare the *dN/dS* not to one (neutral expectation), but to a null expectation under a model of purifying selection (Latrille et al., 2023). We therefore used a mutation-selection model that infers the fitness landscape of a given site in a given proteins, assuming this fitness landscape is shared across mammals (no adaptation). We can then derive from this fitness landscape the expected *dN/dS* at this site, which is usually lower than one (Latrille et al., 2023). If the observed *dN/dS* at the site is significantly higher than the null expectation under purifying selection (*ω*_0_), we can conclude that this site has undergone one or several episodes of positive selection (Latrille et al., 2023). We can also compute a “non-adaptive *dN/dS*” of a protein by taking the average of the *dN/dS* of sites compatible with a model of strictly purifying selection.

To pinpoint which part of the recombination pathway may be subject to different selective pressures, meiotic recombination proteins have been divided into the six steps that follow (Hunter, 2015). 1) DSB formation: recombination initiates by programmed double-strand breaks (DSBs) induced by the protein complex SPO11-TOPVIBL (Keeney et al., 1997; Robert et al., 2016). 2) DSB processing: The hanging ends of the double-stranded DNA are then resected by the MRE11-RAD50-NBS1 complex, leaving DNA single stranded for several hundred base pairs on both sides of the cut (Kim et al., 2025). 3) Strand invasion: Several proteins then stabilize the single stranded ends and allows the formation of a nucleoprotein filament that performs the homology search and the invasion on an intact double-stranded DNA, resulting in chimeric double stranded DNA with one strand from each parental chromosome. 4) CO/NCO decision: Several proteins direct the processing of the recombination intermediate into a reciprocal exchange of chromosome arm (a CO), or a local gene conversion event (NCO). 5) Resolution: Finally, COs and NCOs are resolved and homologous chromosome will segregate in different cells at the end of division I. Additionally, the molecular process of recombination is temporally and functionally linked with the formation of meiosis-specific chromosome axes, the alignment of homologous chromosomes and the formation of the synaptonemal complex that associate intimately the homologous chromosomes on their entire length (step 6, Pairing and synapsis) (Zickler and Kleckner, 2023).

We first confirm that the average *dN/dS* of recombination proteins is significantly higher than that of control proteins (Figure 1A). This is the case both for meiosis-specific and those that also have activities outside of meiosis (pleiotropic proteins), although meiosis-specific tend to have higher *dN/dS* than pleiotropic ones (Figure 1A). Interestingly, the *dN/dS* of proteins involved in different steps of the recombination process do not differ significantly (Figure 1B).

**Figure 1.**
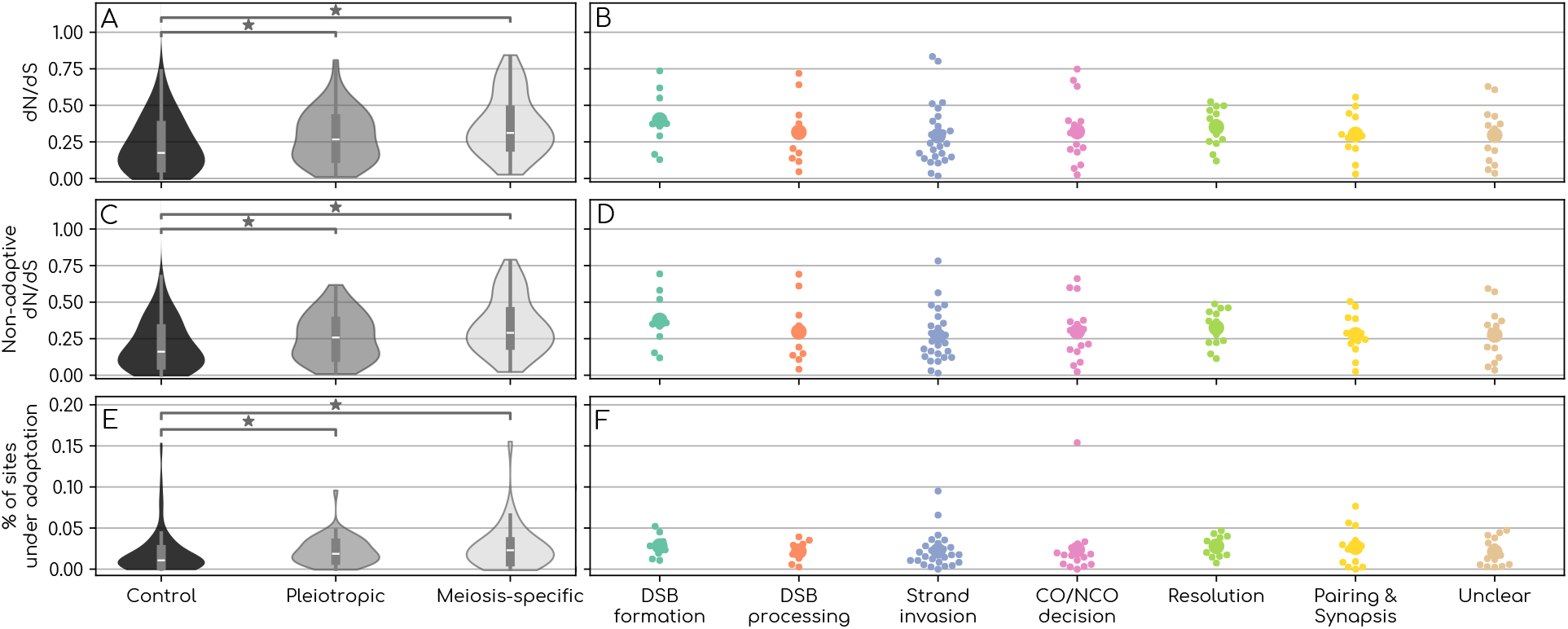
Rate of evolution and proportion of positively selected sites in recombination and control proteins. A,B: average *dN/dS* distribution across recombination and control proteins. C,D: “Non-adaptive” *dN/dS*, averaged across sites with no evidence of positive selection. E,F: Proportion of sites for which the *dN/dS* is higher than under a null expectation of purifying selection. A,C,E: Black violins represent the distributions for randomly selected control proteins, dark gray violins the distribution for genes with functions outside of meiosis, and light gray violins the distribution for meiosis-specific genes. B,D,F: Each dot represents a protein, and large dots represent the mean. We tested for differences between the mean of distributions with a Kruskal-Wallis test. We corrected for multiple tests for all pairs of distributions within a panel with a Holm-šídák test. We represented significant differences at the 95% threshold after multiple test correction with a bar and a star.

Second, proteins involved in meiotic recombination have on average a slightly higher incidence of positive selection compared to control proteins, and this is true both for pleiotropic and meiosis-specific proteins (Figure 1E). Again, the proportion of positively selected sites in proteins involved in different steps of the recombination process do not differ significantly (Figure 1F). However, the contribution of positively selected sites to the *dN/dS* is not significantly different between control, pleiotropic and meiosis-specific proteins (Figure S1A). When comparing the *dN/dS* of these proteins after removing positively selected sites, the *dN/dS* decreases by about 7% in every protein category (Figure 1C & Figure S1A) and this does not change the pattern described in the previous paragraph (Figure 1C&D).

Overall, even if there is significantly more sites under positive selection in recombination proteins compared to control proteins, the increase of the *dN/dS* in these proteins appears to be mainly due to lower purifying selection.

While the analyses presented above inform us on the selection regime of recombination proteins, the biological processes that cause higher positive selection or lower selective pressure cannot be identified by only looking at the average *dN/dS* or the rate of positive selection across mammals. To this end it is more informative to investigate the variation and covariation of recombination protein’s *dN/dS* between mammals. Identifying the factors associated with the evolution of recombination protein’s *dN/dS* in mammals could shed light on the processes that impose selection on the mammalian recombination pathway.

### Explaining variation in selective pressures exerted on recombination proteins

Some proteins evolve quickly in all mammals because they have flatter fitness landscapes. These landscapes can be shared between species, regardless of karyotype, recombination rates, or the specific environment in which the mammals live. This leads to a high *dN/dS* but fairly constant between species. On the other hand, some proteins can have a smaller *dN/dS* on average, because they generally evolve under a constrained fitness landscape, but one that can change depending on the environment. This leads to a lower *dN/dS* on average but that varies significantly between species.

We used a Bayesian approach (Lartillot and Poujol, 2011; Latrille et al., 2021) to estimate variations in *dN/dS* at the protein level along the tree of mammals for both recombination and control proteins. As an illustration, ANKRD31, a gene involved in DSB formation between X and Y chromosomes in males (Boekhout et al., 2019; Papanikos et al., 2019), evolves at a high rate on average (*dN/dS* = 0.75) but with little variation between species (between 0.61 in the gracile shrew mole *Uropsilus gracilis* and 0.82 in the desert woodrat *Neotoma lepida*, Figure S2). On the other hand, HORMAD1, a gene mainly involved in chromosome pairing and synapsis (Shin et al., 2010; Daniel et al., 2011; Dereli et al., 2024) evolves at a lower rate on average (*dN/dS* = 0.41) but one that varies strongly between species (between 0.08 in Hoffmann’s two-toed sloth *Choloepus hoffmani* and 1.57 in the great roundleaf bat *Hipposideros armiger*, Figure S2).

For each of the control and recombination proteins, we can compute the variance in the *dN/dS* across mammals. Both pleiotropic and meiosis-specific proteins have lower variances in *dN/dS* than control proteins, with no significant differences between them (Figure 2A). This indicates that recombination proteins evolve under more stable selective pressures than control proteins. Of the recombination proteins, only those involved in chromosome pairing and synapsis have a higher *dN/dS* variance than those involved in DSB formation, strand invasion and resolution (Figure 2B).

**Figure 2.**
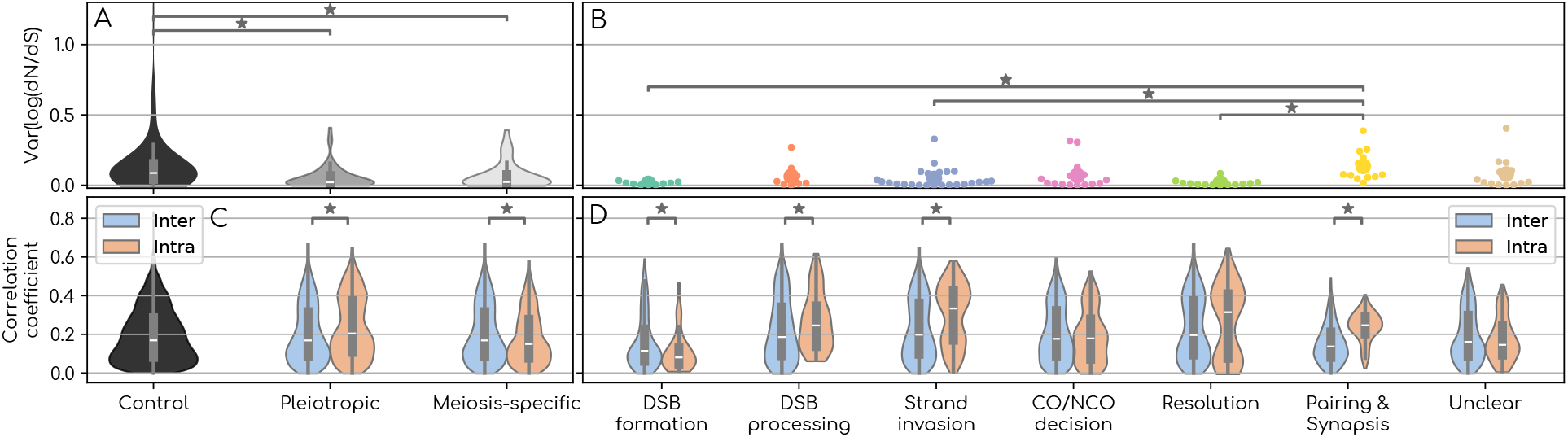
A,B: Distribution of the variance of the log(*dN/dS*) for recombination and control proteins. A: Black violins represent the distributions for randomly selected control proteins, dark gray violins the distribution for genes with functions outside of meiosis, and light gray violins the distribution for meiosis-specific genes. B: Each dot represents a protein, and large dots represent the mean. We tested for differences between the mean of distributions with a Kruskal-Wallis test. C,D: Distribution of correlation coefficients between the log(*dN/dS*) of pairs of proteins. In black we represented the distribution of the correlation coefficient between all pairs of randomly selected control proteins. In orange, we represented the distribution of correlation coefficient between all pairs of proteins belonging to the same category. In blue, for each category, we represented the distribution of correlation coefficients between proteins from this category and proteins outside the category. We tested for differences between pairs of blue and orange distribution with a Kruskal-Wallis test. We corrected for multiple tests within a panel using a Holm-šídák test. We represented significant differences at the 95% threshold after multiple test correction with a bar and a star.

To assess whether this variation reflects functional changes in the recombination process, we tested whether the *dN/dS* of recombination proteins involved in the same steps of the process tend to co-evolve. We therefore inferred the full correlation matrix of the *dN/dS* of the 200 proteins using a newly developed method of phylogenetic and probabilistic principal component analysis (P3CA) (Montoya et al., 2026). This is similar to an evolutionary rate covariation (ERC) analysis (Clark et al., 2013), except that rather than assessing the pairwise correlation between a normalized *dN* between proteins, we estimate the full correlation matrix between *dN/dS* taking into account non-independence between branches due to phylogenetic relatedness. For DSB processing, strand invasion and synapsis, proteins involved in the same step tend to show a greater signal of coevolution among themselves than with proteins involved in other steps of the pathway (Figure 2D). This is striking for proteins involved in synapsis, for which there is almost a two-fold increase in average correlation coefficient (Figure 2D). This suggests that these shifts in *dN/dS* likely reflect functional changes in the recombination process in mammals. On the other hand, for proteins involved in DSB formation, the pattern is reversed but the difference is very small, and may be due to the very low variance in the *dN/dS* of DSB formation proteins in general (Figure 2B&D). It is worth noting that recombination proteins do not show a greater correlation in *dN/dS* among themselves compared to control proteins (Figure 2C).

To identify the biological processes driving this (co)variation, we applied the P3CA separately for recombination and control proteins. Briefly, the P3CA models the evolution of coevolving proteins across the branches of the mammals’ tree in a reduced set of latent variables or “processes” that can for instance capture the various steps of the recombination pathway. This is particularly useful for assessing the relationships across a large number of *dN/dS* variables (see details in the material and methods). These latent variables are reconstructed along the phylogeny, and the extent to which they explain the *dN/dS* evolution of the different proteins is given by their loadings (a matrix correlating each protein to each latent variable). A protein with a positive loading on a given latent variable has a positive correlation between its *dN/dS* and the latent variable and a negative one with a negative loading. We estimated the number of unobserved or latent variables using cross-validation (details in Material and Methods).

Only three independent latent variables are necessary to explain the observed *dN/dS* covariation patterns for recombination proteins, and two for control proteins (Figure S4). The rest of the variance is best explained by an independent random noise, that may reflect either estimation errors of the *dN/dS*, or protein-specific dN/dS evolution along the tree (see details in Material and Methods). The first latent variable of recombination proteins (PC1) explains 38.4% of the total variance in *dN/dS* and is approximately partitionning almost all the recombination proteins into two groups of interrelated proteins. The first group comprises proteins that are primarily involved in chromosome pairing and synapsis, and exhibit an increase in *dN/dS* with PC1. The second group comprises almost all the other proteins, which tend to decrease in *dN/dS* with PC1 (Figure 3C). For the second latent variable (PC2), which explains 7.3% of the variance, all proteins that have a loading on it have a negative one, meaning that they tend to increase in *dN/dS* when PC2 decreases (Figure 3C). Proteins involved in chromosome pairing and synapsis are also over-represented on this axis. The last latent variable (PC3) only explains 2% of the total variance, and is mostly showing opposite direction of evolution between RPA2 (positive loading) and TEX11 (negative loading; see Figure 3C). Of note, adding this axis to the two previous ones does not improve much the fit of the model (Figure S4), and therefore its functional relevance is unclear.

**Figure 3.**
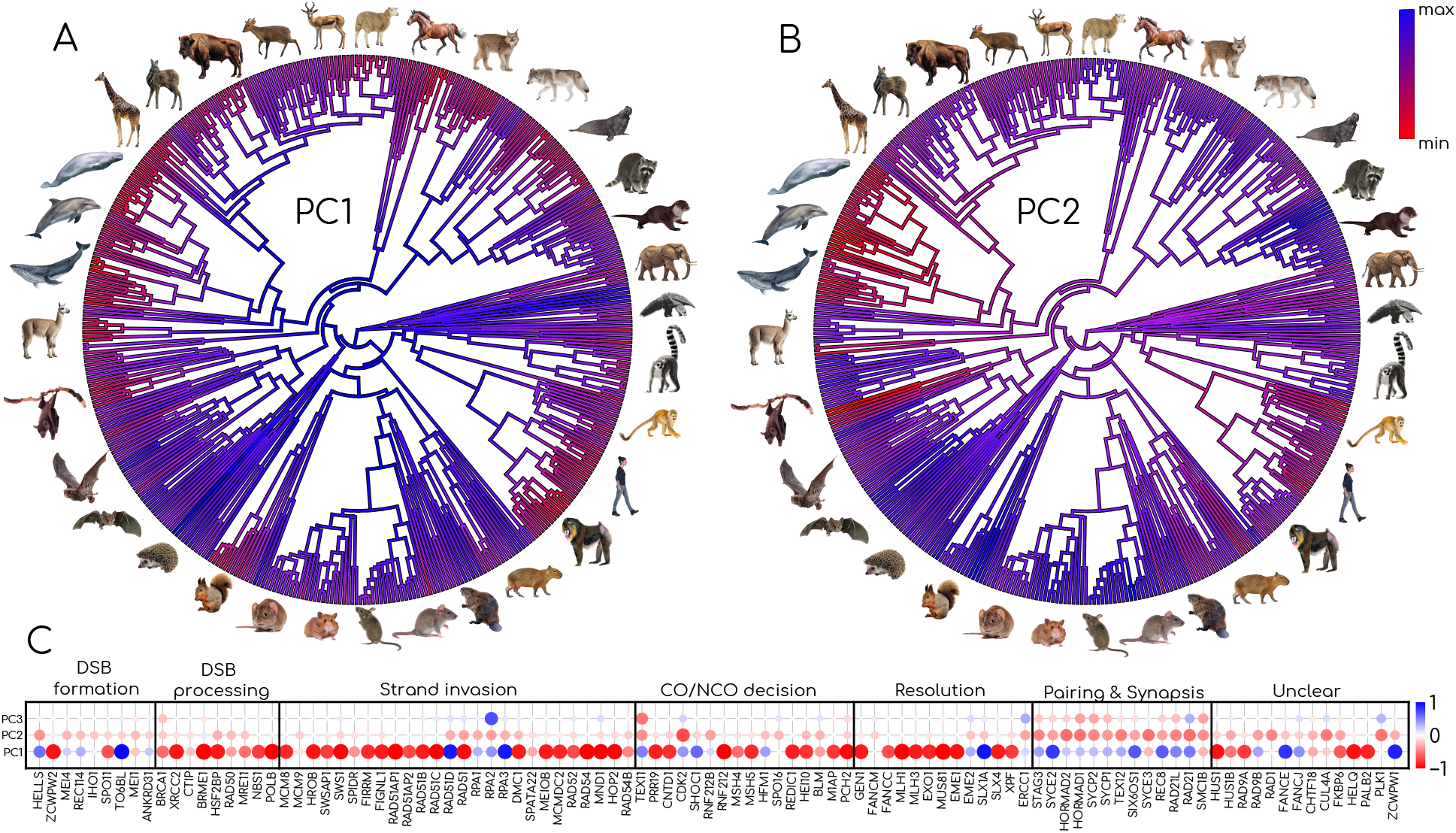
A,B: Evolution of PC1 and PC2 along the phylogeny of placental mammals. PC1 and PC2 are expressed in arbitrary units. C: Loadings between recombination proteins and the PC axes. Each dot represents a protein, the size gives the strength of the correlation between the protein’s *dN/dS* and the PC axis, and the color gives the direction. Along a given axis, blue proteins have a higher *dN/dS* in blue lineages, and red proteins have a higher *dN/dS* in red lineages.

By looking at how PC1 unfolds over the phylogeny, one striking pattern is its evolution with time (Figure 3A). The value of PC1 is very high at the root of the phylogeny and generally decreases towards the tips of the tree (Figure 3A). Therefore, it correlates with proteins whose *dN/dS* evolve with a temporal trend. Proteins with a negative loading on PC1 tend to have increasing *dN/dS* (decreasing selective pressures) with time (e.g. FIGLN1) (Figure 4A). On the other hand proteins with a positive loading on PC1 tend to have decreasing *dN/dS* (increasing selective pressures) with time (e.g. SYCE2) (Figure 4A). However, the value of PC1 does not increase at the same rate in different species. To understand which biological processes PC1 reflects, we compiled life-history traits, karyotypic and genome-wide recombination rate data available from existing databases, and tested whether some of these traits were also related to PC1.

**Figure 4.**
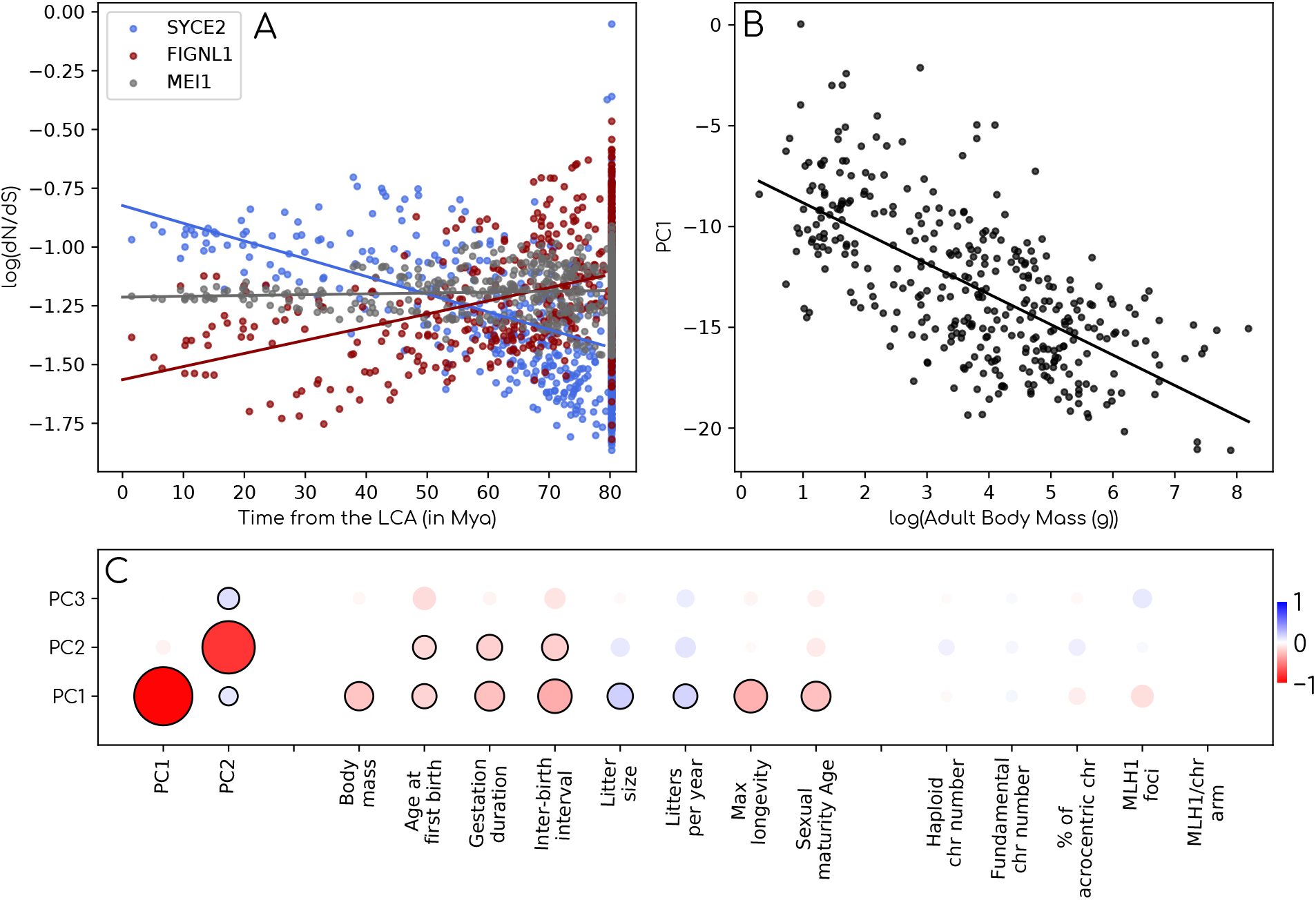
A: Evolution of the log(dN/dS) as a function of time for three recombination proteins. B: Linear regression between PC1 and adult body mass in placental mammals. C: Phylogeny-corrected pairwise correlations between PC axes of recombination proteins, PC axes of control proteins and different traits associated to the slow-fast continuum or karyotypic/recombination traits. Pairwise correlations were computed using a Pagel’s lambda model implemented in *phylolm*. The size of dots represent the strength of the correlation, and the color the direction. Significant correlations at the 95% threshold are surrounded by a black circle.

Interestingly, neither karyotypic characteristics such as haploid or fundamental chromosome number or recombination rate per meiosis or per chromosome arm are significantly correlated to PC1 (Figure 4C). On the other hand PC1 is significantly correlated with all traits related to the slow-fast continuum in mammals (Figure 4B&C). The *dN/dS* of proteins that have a negative loading on PC1 (red dot in Figure 3) is generally higher in species with a slow pace of life (red lineages in Figure 3A), while the *dN/dS* of proteins that have a positive loading on PC1 (blue dots in Figure 3C) is generally higher in species with a fast pace of life (blue lineages in Figure 3). Interestingly, PC1 is almost perfectly correlated to PC1 of control proteins (Figure 4C). This suggests that PC1 reflects factors influencing *dN/dS* evolution in the whole proteome.

PC2 does not have a strong temporal trend (Figure 3B), and does not correlate either with karyotype or genome-wide recombination rate (Figure 4C). However, it also correlates (more weakly) with some pace of life-associated traits (age of the female at first birth, gestation duration and inter-birth interval) (Figure 4C). Again, the *dN/dS* of proteins that have a negative loading on PC2 is generally higher in species with a slow pace of life. This latent variable is also highly correlated to the second latent variable of control proteins (Figure 4C), again suggesting that it influences *dN/dS* evolution in the whole proteome.

While chromosome number and recombination rate may not be the main drivers of the evolution of the recombination pathway, they could still influence the *dN/dS* of isolated proteins. We therefore looked at the pairwise correlation between each of the 200 proteins’ *dN/dS* and haploid chromosome number, fundamental chromosome number, recombination rate per meiosis, or per chromosome arm. After correcting for multiple testing, none of the proteins had a significant correlation between their *dN/dS* and karyotypic/recombination traits. Moreover, correlation coefficients and p-values are similar between recombination and control proteins (Figure S5).

## Discussion

In this study, we sought to explain why recombination proteins evolve at a high rate in mammals, and what ecological/demographic/cellular processes could explain variation in their selective constraints. We showed that the evolutionary rate of recombination proteins tend to be higher for meiosis-specific proteins. This is consistent with studies showing that meiosis-specific genes evolve at a higher rate than their mitotic paralog in mammals (Boekhout et al., 2019). We show that this rapid evolution can be explained by reduced purifying selection with a more limited role of positive selection. We also showed that these selective pressures are generally conserved between mammals. We identified the major axes of variation in evolutionary rates of recombination proteins, and contrary to previous assumptions, we showed that these axes were largely unrelated to the evolution of the number/type of chromosomes or to genome-wide recombination rates.

### On the selective pressures exerted on the mammalian recombination pathway

We showed that the variation in evolutionary rate of the recombination pathway can be summarized by three latent variable. The first latent variable, PC1, both evolves in a directional way with time, and across extant species, correlates with pace of life-associated traits. There are three main hypotheses to explain the existence of this first axis. First, *dN/dS* in terminal branches can be overestimated because segregating deleterious polymorphisms that will never reach fixation can artificially increase the *dN/dS* in short terminal branches, but will barely impact internal branches (Mugal et al., 2014, 2020). Second, residual alignment errors will usually inflate the *dN/dS* especially in terminal branches (Jordan and Goldman, 2012; Selberg et al., 2025). These two explanations predict an increase in *dN/dS* in recent times, but do not explain the correlation with pace of life-associated traits. The last hypothesis is a survival bias argument. Those lineages that gave rise to extant species are the ones that kept a population size large enough such that they did not go extinct. This induces a temporal bias, whereby lineages that are deeper in the phylogeny have evolved under higher effective population size. Because of larger population sizes, these lineages are more likely to have higher efficacy of natural selection, and therefore a lower *dN/dS*. This last argument predicts both a gradual increase in *dN/dS*, and an increase in species with small population sizes (more often slow-living species). However, under all those hypotheses, all *dN/dS* should correlate positively with PC1 and evolve in the same direction. Instead, many genes decrease in *dN/dS* with time, and for meiotic genes, they tend to be functionally related, another observation not predicted by the three scenario presented above.

Interestingly, all control and recombination proteins have a positive loading on PC2. This suggests that PC2 represents a species/node effect, whereby the *dN/dS* of all genes correlate in the same direction. Moreover, PC2 also correlates with pace of life-associated traits, independently from PC1. Thus, PC2 is more likely to capture an effective population size effect than PC1. This implies that, in both control and recombination proteins, effective population size is not what explains most of the variation in protein’s *dN/dS*. Instead, other processes linked to the slow-fast continuum largely affect the rate of protein evolution along the genome. Some proteins tend to evolve more slowly in slow-living species than in fast-living species, and the reverse for others.

### The genetic basis of recombination rate evolution in mammals

Previous molecular evolution studies on the mammalian recombination pathway were conducted to shed light on the genetic basis of recombination rate evolution. In one study, the authors found one gene (TEX11) whose *dN/dS* is associated with genome-wide recombination rates (Dapper and Payseur, 2019). With our dataset, we do not recover this correlation. This correlation was previously found using recombination data for 9 species, but was not significant when correcting for multiple testing (Dapper and Payseur, 2019). After applying multiple-test correction to a larger sample (36 species), we could not find any protein whose *dN/dS* was significantly associated with recombination rates. Therefore, it seems that the association between recombination rates and the rate of evolution of recombination proteins is only minor. This is particularly surprising because in several mammalian species, variation in recombination rates within population is clearly and repeatedly associated to large effect loci, close or within some of the proteins examined in this study (Johnston, 2024; Brekke et al., 2025). There are three possible reasons for this discrepancy. First the causal mutations that affect recombination rates could mostly appear in regulatory regions, and change the timing of expression or the abundance of recombination proteins without affecting their coding sequences. Second, is it possible that new mutations modifying recombination rates are neutral or deleterious and rarely reach fixation. Indeed, the fact that recombination rate is variable and has a genetic basis does not imply that this variation is adaptive (Payseur, 2025). The higher or lower recombination rates experienced by some individuals compared to the common ancestor may have no significant effect on fitness, or may be deleterious. Finally, it is possible that a few large-effect mutations in coding sequences do allow adaptive changes in recombination rates but that most non-synonymous substitutions in recombination proteins are related to other processes, as developed below.

### Longevity and synapsis proteins

Interestingly, when we consider the *dN/dS* variation explained by the first latent variable, we find that proteins involved in chromosome pairing and synapsis (all except SYCE3 and SMC1-*β*) tend to evolve at a slower rate in long-lived species. In female mammals, prophase I starts in the embryo and is blocked with all homologous chromosome paired until fecundation. In species with long lifespan and old age of sexual maturity (e.g. primates or cetaceans) these chromosomes have to stay paired for decades, through the maintenance of sister chromatid cohesion. In mice, defects in cohesins have been associated with increased aneuploidy with maternal age (Prieto et al., 2004; Revenkova et al., 2004; Hodges et al., 2005; Lister et al., 2010; Leem et al., 2025). In human females, age-related deterioration of meiotic cohesin (such as REC8, RAD21, RAD21L) and its protection is widely regarded as a major driver of oocyte aneuploidy, contributing substantially to infertility, miscarriage, and adverse pregnancy outcomes (Chiang et al., 2010; Duncan et al., 2012; Nagaoka et al., 2012; Mihalas et al., 2024). It appears probable that the stability of proteins responsible for the maintenance of pairing is more critical for fertility in long-lived species, and therefore slightly destabilizing non-synonymous mutations are less tolerated. This would explain why the rates of evolution of cohesins, and of the associated structural components of the meiotic chromosome axes and the synaptonemal complex, tend to decrease in long-lived taxa. Importantly, the variance decomposition approach taken here was essential to reveal this pattern. Indeed, the intensification of selection on pairing and synapsis proteins (and many others) in long-lived species is masked by the overall lower selection efficacy in those species (Jobson et al., 2010), probably due to small population size (Brevet and Lartillot, 2021; Latrille et al., 2021). Decomposing the variance shared by different proteins into different latent variables appears to be an effective strategy to decouple the slow-fast effect from the effect of the effective population size on *dN/dS* evolution.

### Conclusion

Our study reveals that the rapid evolution of mammalian meiotic recombination protein is largely driven by globally reduced levels of purifying selection compared to the rest of the proteome. However, we show that there is variation in the selective pressures exerted on recombination proteins in mammals, which is poorly explained by genome-wide recombination rates or chromosome number. Instead, this variation appears to reflect differing constraints on chromosome pairing and synapsis across species with contrasting lifespans. In particular, long-lived species may experience stronger selective constraint due to the difficulty of maintaining chromosome pairing over decades in female oocytes. These findings highlight that divergence in recombination machinery can be shaped by life-history–dependent constraints that are largely independent of recombination rate and its effects on genetic diversity or selection efficacy.

## Material and Methods

### Alignments and Phylogenetic Tree

From the literature, we identified 114 genes with an experimentally validated role during meiotic recombination in mouse. We searched for multiple species alignments of these proteins in the TOGA ortholog database (Kirilenko et al., 2023). These orthologs have been annotated with a machine-learning methods based on whole genome alignments (Kirilenko et al., 2023). They were subsequently aligned with MACSE2 (Ranwez et al., 2018) in Kirilenko et al. (2023). We found alignments for 100 of the recombination proteins, listed in Supplementary Data S1. Among the 17,339 alignments remaining, we randomly drawn 100 as a control dataset. The mammalian tree used throughout the study was generated in the study of Álvarez Carretero et al. (2022) integrating genomic data from 72 species and genetic markers for 4705 species.

### Alignment cleaning

We first removed sequences from species that were absent in the tree from the alignments. When the alignment contained multiple sequences from the same species, we only kept sequences coming from the assembly with the highest quality (highest Scaffold N50). After this step, the alignments still contained some paralogs. We therefore built gene trees for each of the alignments with a GTR+G4 model using the iqtree software (Minh et al., 2020). When multiple paralogs were monophyletic, we built a consensus sequence by replacing variable nucleotides by a N, and removed all paralogs when paraphyletic. We then used HMMcleaner (Di Franco et al., 2019) to remove misaligned portions in all alignments. We then used PhylteR (Comte et al., 2023) with default parameters to remove outlier sequences, whose phylogenetic placement significantly departs from the rest of the sequences of the species. This procedure usually removes mis-annotated sequences and hidden paralogs, but do not affect small phylogenetic discrepancies such as the ones caused by incomplete lineage sorting (Comte et al., 2023). Despite all these curation steps, some residual alignment errors remained. We therefore visually inspected and manually curated all the alignments. For each protein, any residual misalignment for each species were reported in a file. Any species with more than 5% of the alignments with at least one residual error were removed from all alignments (12 species).

### Evolutionary rates estimations

To estimate site-wise *dN/dS* across mammals, we used a Bayesian version of the Muse and Gault (Muse and Gaut, 1994) codon model implemented in Latrille et al. (2023) with default settings. This method uses an MSA and a phylogenetic tree to reconstruct the posterior distribution of the *dN/dS*. We then used a mutation-selection model to estimate the posterior distribution of the expected *dN/dS* at each site under a null model of purifying selection (see Latrille et al. (2023) for more details on the model). We classified a site as being under positive selection if the 95% credibility interval of its *dN/dS* was above and not overlapping the 95% credibility interval of its “null” *dN/dS* (*ω*_0_). To calculate the average *dN/dS* of a protein after accounting for the effects of positive selection, we averaged the *dN/dS* across sites that were not classified as undergoing positive selection.

To investigate variations in the *dN/dS* over the phylogeny of mammals, we used *nodeomega* (Latrille et al., 2021), which is a more efficient re-implementation of the *coevol* model (Lartillot and Poujol, 2011). This method estimates the posterior distribution of the average *dN/dS* over a protein at each node of a given tree, by using the information of substitution rates in the surrounding branches. Importantly, with maximum likelihood estimation, small branches with few substitutions often lead to extreme *dN/dS* values in the surrounding nodes, leading to incorrect estimations of the variance of *dN/dS*. The method *nodeomega* deals with this issue by imposing a gaussian prior to the *dN/dS* of each node, centered on the value of the parent node. This way, the *dN/dS* varies in the next nodes only if there is enough signal in the substitutions that support a higher or lower *dN/dS* compared to the previous node. we ran *nodeomega* with default parameters, using 1200 iterations and a burn in of 200. For each node, we considered the median of the posterior distribution of *dN/dS*.

### Phylogenetic probabilist principal component analysis

To estimate co-variation in the *dN/dS* of recombination and control proteins, we used a phylogenetic probabilistic principal component analysis (P3CA ; Montoya et al. (2026)).

The P3CA model assumes that a small but unknown (*q*) number of independent univariate processes (*z* variables, hereafter referred to as “latent variables”) can be used to model a larger number (*p*) of interdependent *dN/dS* variables evolving along the branches of the mammal tree. Therefore, the model offers a more parsimonious description of coevolving proteins. More formally, the model is described by the following equation:

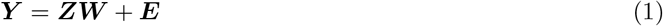

***Y*** is a matrix containing the standardized and centered *dN/dS* of *p* proteins for *n* species (size *n* × *p*); ***Z*** is an *n* × *q* matrix containing *q* latent variables; and ***W*** is a *q* × *p* matrix relating the latent variables to the observed variables. The *q* latent variables follow a Gaussian process, *z* ∼ *N* (0, ***C***). Where ***C*** is the *n* × *n* variance covariance matrix of a Brownian motion process unfolding on the tree (see Felsenstein 1973). Finally, ***E*** is an n by p matrix containing p independent vectors of phylogenetically correlated noise, *e* ∼ *N* (0, *σ*^2^***C***). The model’s parameters (***W*** and *σ*^2^), including the latent variables (***Z*)**, are inferred using an Expectation-Maximization (EM) algorithm. This enables estimation of the P3CA and imputation of missing values in ***Y*** (i.e. where some proteins are missing for certain species).

From ***W*** and *σ*^2^, we can compute the covariance matrix (***R***) between the *dN/dS* of all pairs of proteins:

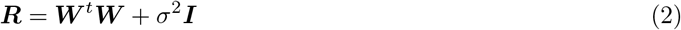

As the number (*q*) of latent variables is unknown, it was estimated using cross-validation. This procedure avoids overfitting due to the inclusion of too many latent variables and missing meaningful covariation by including too few latent variables to appropriately describe co-variation between proteins. Cross-validation was achieved by randomly removing 1% of the entries in ***Y*** before running the P3CA to estimate these missing values across a range of latent variables (we varied the number of latent variables between 2 and 50). For each number of latent variables, we repeated the procedure 100 times and calculated the average mean-square error between the true values of ***Y*** (i.e. the values that were removed) and the inferred values. We selected the number of variables that best predicted the missing entries (see Figure S4).

### Life history traits

We obtained the life-history traits associated with the pace of life from the Pantheria database (Jones et al., 2009). We used karyotype information compiled in the study of Blackmon et al. (2019), available for 222 species from our dataset, and genome-wide recombination rates compiled in Rossi and Pigozzi (2025), available for 36 species from our dataset. We regressed these traits and either protein’s *dN/dS* or P3CA latent variables using a PGLS with Pagel’s lambda model (Revell, 2010; Clavel and Morlon, 2020) implemented in *phylolm* (Tung Ho and Ané, 2014).

## Data availability

All data associated with this study were deposited at https://doi.org/10.5281/zenodo.20489001. All codes necessary to reproduce this study are available at https://gitlab.in2p3.fr/julien.joseph/meionet.

## Acknowledgments

We wish to thank Philippe Veber for insightful discussions and Thibault Latrille for helpful comments on a previous version of this manuscript. This work was performed using the computing facilities of the CC LBBE/PRABI.

