## Supplementary Figures for "Explaining the rapid evolution of mammalian meiotic recombination proteins"

### Supplementary material from: Explaining the rapid evolution of mammalian meiotic recombination proteins

April 28, 2026

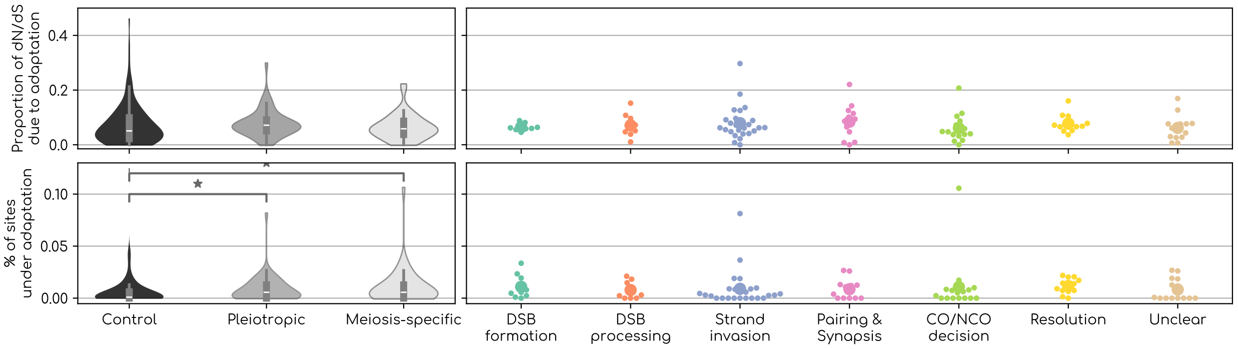


Figure S1: A,B: Proportion of the $dN/dS$ explained by sites under adaptation. It can be computed by comparing the total $dN/dS$ to the one after removing sites under adaptation ($dN/dS > \omega_0$. C,D: Proportion of sites with a $dN/dS > 1$ A,C: Black violins represent the distributions for randomly selected control proteins, dark gray violins the distribution for genes with functions outside of meiosis, and light gray violins the distribution for meiosis-specific genes. B,D: Each dot represents a protein. We tested for differences between the mean of distributions with a Kruskal-Wallis test. We corrected for multiple tests for all pairs of distributions within a panel with a Holm-Šídák test. We represented significant differences at the 95% threshold after multiple test correction with a bar and a star.

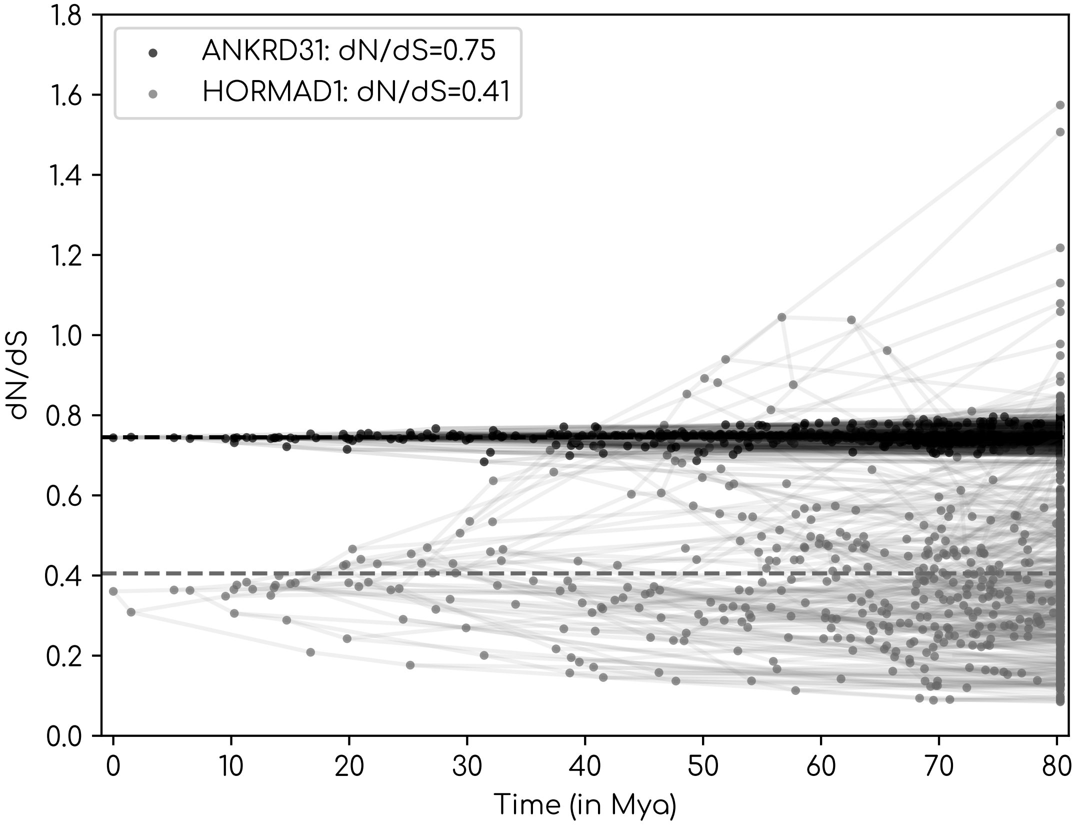


Figure S2: Evolution of the $dN/dS$ of ANKRD31 (black) and HORMAD1 (grey) in mammals. Dots represents node of the tree and edges represent branches. $dN/dS$ at each node were reconstructed using the medians of the posterior distributions of *nodeomega* (Latrille et al., 2021)

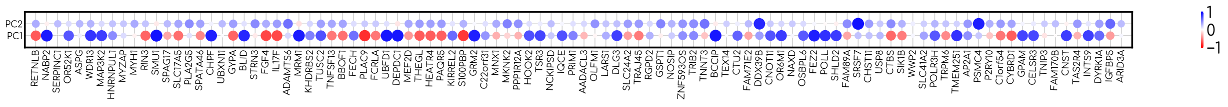


Figure S3: Loadings between control proteins and the PC axes. Each dot represents a gene, the size gives the strength of the correlation between the protein's $dN/dS$ and the PC axis, and the color gives the direction. Along a given axis, blue proteins have a higher $dN/dS$ in blue lineages, and red proteins have a higher $dN/dS$ in red lineages.

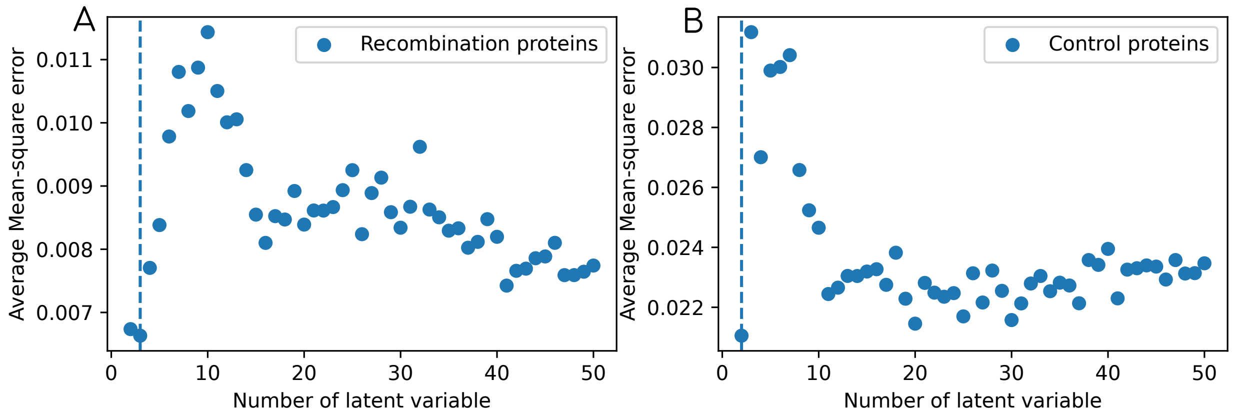


Figure S4: Average mean-square errors as a function of the number of latent variable for recombination proteins (A) and control proteins (B). We removed from the matrix $\boldsymbol{Y}$ 1% of the data, and ran the P3CA several times. We varied the number of latent variable between 2 and 50, and repeated the procedure 100 times for each number of latent variable. For each number of latent variable, we plotted in blue at the averaged mean-square error between the true and the inferred value of the removed entries of $\boldsymbol{Y}$ across the 100 replicates. The vertical dashed blue line represents the number of variable that best predict those missing entries (3 for recombination proteins and 2 for control ones).

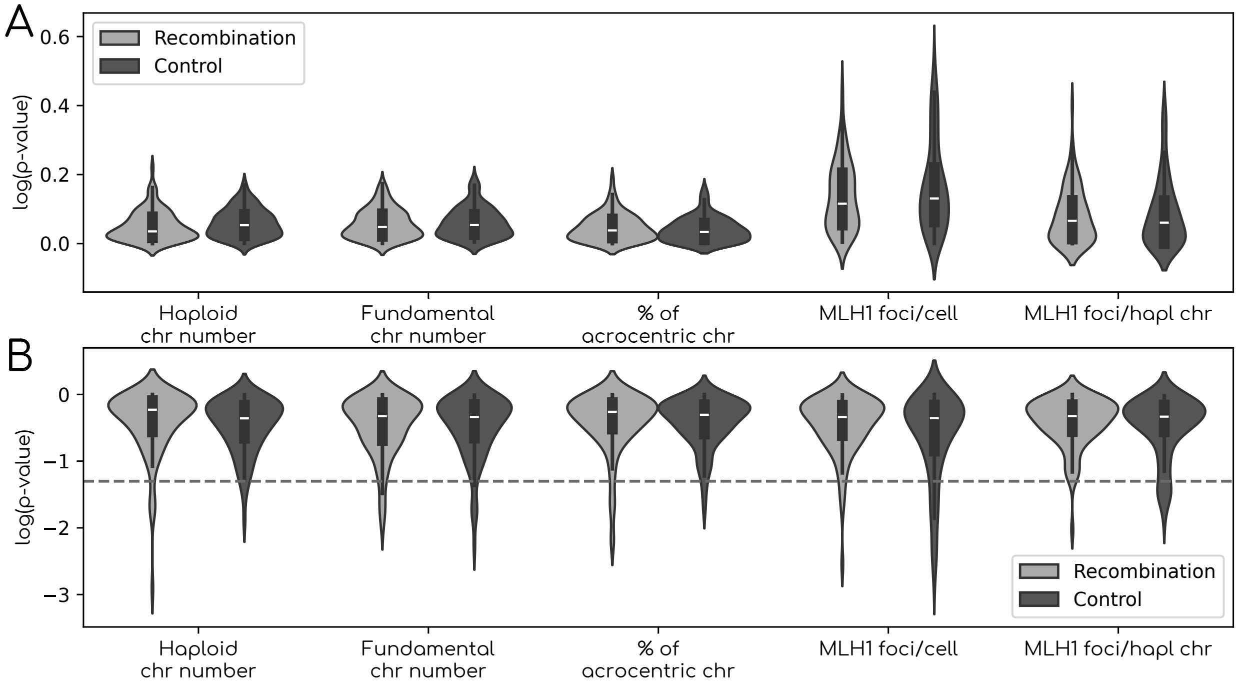


Figure S5: Distribution of orrelation coefficients between the $dN/dS$ of protein and karyotipic/recombination traits (A) and associated p-values (B). Correlations were tested using a Pagel's lambda model (**ref**) implemented in *phylolm* (Tung Ho and Ané, 2014).
